# Her9 is required for the migration, differentiation, and survival of neural crest cells

**DOI:** 10.1101/2025.07.18.665589

**Authors:** Cagney E. Coomer, Sumanth Manohar, Evelyn M. Turnbaugh, Ann C. Morris

## Abstract

Neural crest cells (NCC) are vertebrate-specific multipotent progenitor cells that arise from the neural plate border and go on to contribute to a wide variety of morphological structures such as the jaw and palate, enteric nervous system (ENS), and pigment cells. Defects in essential steps in neural crest cell development have been associated with a wide variety of congenital disorders, collectively referred to as neurocristopathies. Her9/Hes4 is a bHLH-O transcriptional repressor that has been shown to regulate neural crest cell and craniofacial development in Xenopus and zebrafish, however the extent of Her9 function in other neural crest cell lineages has not been investigated. In this study, we characterized NCC phenotypes in *her9* mutant zebrafish. We show that loss of Her9 perturbs the development of several NCC derivatives. *Her9* mutants display a variety of NCC defects, including craniofacial abnormalities, alterations in pigment cell lineages, and improper formation of the gut. These phenotypes are associated with defects in neural crest cell specification, migration, and differentiation, as well as an upregulation in expression of BMP ligand genes. Furthermore, loss of Her9 leads to apoptosis of NCC derivatives. Collectively, our results show that Her9 functions in neural crest development by regulating members of the NCC gene regulatory network (GRN) to control NCC specification, migration, differentiation and survival.

## Introduction

Neural crest cells (NCC) are a population of vertebrate-specific, multipotent migratory cells that give rise to a wide variety of cell types, such as craniofacial cartilage, pigment cells, peripheral nerves, and glia (Bronner and Simões-Costa, 2016; Rocha et al., 2020). NCC are divided into subpopulations based on their region of origin; these include the cranial neural crest cells (CNCC; craniofacial skeleton, cranial ganglia and heart muscle), vagal neural crest cells (VNCC; enteric neurons and smooth muscle), and trunk neural crest cells (TNCC; dorsal root and sympathetic ganglia, pigment cells), each of which displays distinct migratory patterns and differentiates into several derivatives. NCC development begins during late gastrulation with their induction and specification at the interface between the neural and non-neural ectoderm on the dorsal side of the neural tube at the neural plate border (Kelsh et al., 1996; Schumacher et al., 2011).

Following induction and specification, NCCs go through an epithelial-to-mesenchymal transition (EMT), during which time they transition to migrating mesenchymal cells by changing their morphology, adhesion properties, polarity, and behavior (Theveneau and Mayor, 2012). NCCs then migrate along conserved pathways for long distances to reach their final destinations, differentiating into a wide variety of cell types (Bronner and Simoes-Costa, 2016). Cranial NCC migration begins when a subset of cranial neural crest cells (CNCC) from the midbrain travels anteriorly between and around the eyes, towards the neurocranium (Knight and Schilling, 2006). CNCCs that are posterior to the midbrain/hindbrain boundary migrate ventrally in streams to form the pharyngeal arches and viscerocranium (Schilling and Kimmel, 1994). In contrast, trunk neural crest cells (TNCC) migrate in two pathways, first between the neural tube and somites, and then laterally between the somites and ectoderm (Raible et al., 1992). The first set of cells to migrate gives rise to sympathetic and sensory neurons, Schwann cells, and pigment cells; the second set produces only pigment cells (Raible and Eisen, 1994).

Once migration is complete and NCCs reach their designated locations, they differentiate into specific derivatives. Chondrocytes originate from CNCC and give rise to a significant portion of the vertebrate craniofacial skeleton, including the pharyngeal arches, the mandible, and most components of the jaw (Takahashi et al., 2001). In zebrafish, pigment cells derived from TNCC differentiate into 3 subpopulations: xanthophores, melanophores and iridophores. NCCs also give rise to several neurogenic derivatives in the cranial region (cranial ganglia, Schwann and satellite cells) and in the trunk (dorsal root ganglia, autonomic sympathetic neurons, and Schwann cells) (Kague et al., 2012; Raible and Eisen, 1994; Raible et al., 1992). The enteric nervous system (ENS) is comprised of vagal neural crest (VNCC), which in zebrafish emerges from the hindbrain adjacent to somites 1-7 (Hutchins et al., 2018). Coined the “second brain”, the ENS is responsible for the innervation of the intestine, which mediates the motility of the gut, secretions, and local blood flow (Ganz et al., 2016).

A complex gene regulatory network involving numerous transcription factors and signaling molecules directs the specification, migration, and differentiation of NCCs (Hovland et al., 2020); components of the neural crest GRN are used reiteratively at several developmental stages. For example, transcription factors required for NCC specification include Sox10, Snail1b, Foxd3 and Twist, among others (Carney et al., 2006; Das and Crump, 2012; Thisse et al., 1995; Wang et al., 2011). In addition to its role in NCC specification, Sox10 is required for the migration of vagal NCCs to the intestine (Elworthy et al., 2005; Kapur, 1999; Lake and Heuckeroth, 2013). And finally, Sox10 is required for the development of individual and overlapping NCC derivatives. For instance, in the zebrafish *colourless* mutant, the loss of Sox10 results in defects in melanoblasts, neurons, and glia, but the cranial NCC derivatives are not affected (Dutton et al., 2001). Decades of studies using several vertebrate model organisms have provided a great deal of insight into neural crest cells and their development, but there is still much to discover about the mechanisms and missing components of the neural crest GRN.

Defects in NCC development at any of the steps described above lead to developmental disorders termed neurocristopathies; some examples of neurocristopathies include Hirschspring disease, CHARGE syndrome, Waardenburg syndrome, and neuroblastoma (Etchevers et al., 2019). Neurocristopathies encompass a wide range of phenotypes, including cranioskeletal defects, vision and hearing problems, digestive system malformations, heart defects, loss of skin pigmentation, and cancer (Frisdal and Trainor, 2014). Given the pleiotropic functions of NCC developmental genes, neurocristopathies rarely involve just one tissue, and often exhibit variable expressivity (Etchevers et al., 2019). For example, Hirschsprung disease (HSCR), which has been linked to mutations in the NCC genes *SOX10*, *PAX3*, *MITF*, and *SNAIL*, is primarily characterized by abnormal neurogenesis in the intestine which leads to blockage of stool. Patients with HSCR also suffer from piebaldism (the congenital absence of melanocytes from areas of the skin) and may have hearing loss (Le Douarin et al., 2004; Steel and Barkway, 1989). To improve treatment and therapeutic approaches for neurocristopathies, a better understanding of the genetic mechanisms underlying NCC development, migration, and differentiation is needed.

Her9/Hes4 (also referred to as *Xhairy2* in Xenopus), is a member of the bHLH-O family of transcriptional repressors and is a presumptive NCC gene. In zebrafish and Xenopus, Her9 is expressed at the neural plate border during the window of NCC induction (Tsuji et al., 2003), and additionally in sorted cranial neural crest cells in zebrafish (Stenzel et al., 2022). In Xenopus, morpholino knockdown of *her9* resulted in the repression of neural crest marker genes such as *slug* and *foxd3*, which in turn affected the proliferation and survival of some neural crest (cranial and pigment) derivatives (Glavic et al., 2004). Furthermore, zebrafish *her9* mutants were reported to display bone mineralization defects of the craniofacial skeleton (Stenzel et al., 2022). In our previous study of the function of Her9 in retinal development, we noted that *her9* mutants also displayed a range of craniofacial defects (Coomer et al., 2020). Taken together, these results are consistent with a critical function for Her9 in CNCC differentiation; however, it was not known whether Her9 plays a wider role in the development of other NCC lineages. To that end, we undertook an in-depth characterization of neural crest cell phenotypes in the zebrafish *her9* mutant. We found that, in addition to craniofacial defects, *her9* mutant zebrafish displayed alterations in pigment cell lineages, with increased xanthophores and reduced melanophores and iridophores compared to wild type (WT) zebrafish. *Her9* mutants also displayed defects in intestinal development, including reduced numbers of enteric neurons and glia, and abnormal passage of food through the gut. Finally, loss of Her9 was associated with an increase in apoptosis in NCC-derived tissues. Collectively, our results suggest that Her9/Hes4 is critical for the development and survival of multiple NCC lineages.

## Materials and Methods

### Zebrafish line maintenance

All zebrafish lines were bred, housed, and maintained at 28.5°C on a 14 hour light: 10-hour dark cycle, in accordance with established protocols for zebrafish husbandry (Westerfield, 2000). The *her9* mutant zebrafish line (1 bp insertion resulting in a frame shift and premature termination) was generated via CRISPR-Cas9 genome targeting and has been previously described (Coomer et al., 2020). The *Tg(Sox10:EGFP)*^ba2^, *Tg(Sox10:mRF*P)^vu234^, and *Tg(foxd3:GFP)*^zf105^ transgenic lines (hereafter called *sox10*:GFP, *sox10*:RFP, and *foxd3*:GFP) have been previously described (Carney et al., 2006; Gilmour et al., 2002; Kucenas et al., 2008; Wolman et al., 2008), and were generously provided by Jakub Famulski (University of Kentucky, Lexington, KY). The *Tg(gfap:GFP)^mi2001^* line (hereafter referred to as *GFAP:GFP)*, was previously described (Bernardos and Raymond, 2006) and was obtained from the Zebrafish International Resource Center (ZIRC, Eugene, OR). Embryos were anesthetized with Ethyl 3-aminobenzoate methanesulfonate salt (MS-222, Tricaine, Sigma-Aldrich, St. Louis, MO) and adults were euthanized by rapid cooling as previously described (Wilson et al., 2016). All animal procedures and experimental protocols were approved and carried out in accordance with guidelines established by the University of Kentucky Institutional Animal Care and Use Committee (IACUC), the University of Kentucky Institutional Biosafety Committee, and the ARVO Statement for the Use of Animals in Ophthalmic and Vision Research.

### Her9 mutant genotyping

Genomic DNA (gDNA) was extracted from whole embryos or from tail clips of adult fish. The embryos or tails were placed in 15 ul of 1x Thermopol buffer (NEB, Ipswich, MA) and 7 ul of Proteinase K (Pro K; Sigma-Aldrich, St. Louis, MO) was added to each and incubated at 55° C overnight. After at least 20 hours of digestion, tubes containing the digested tissue were incubated at 95° C for 15 min to deactivate the Pro K. The gDNA was then amplified using *her9* specific PCR primers (**Table 1**) (Coomer et al., 2020). The *her9* WT and mutant alleles were identified by RFLP of PCR products using the BfaI restriction enzyme (NEB: R0568S) as described (Coomer et al., 2020). The restriction digests were visualized on a 2% agarose gel.

**Table 1.**
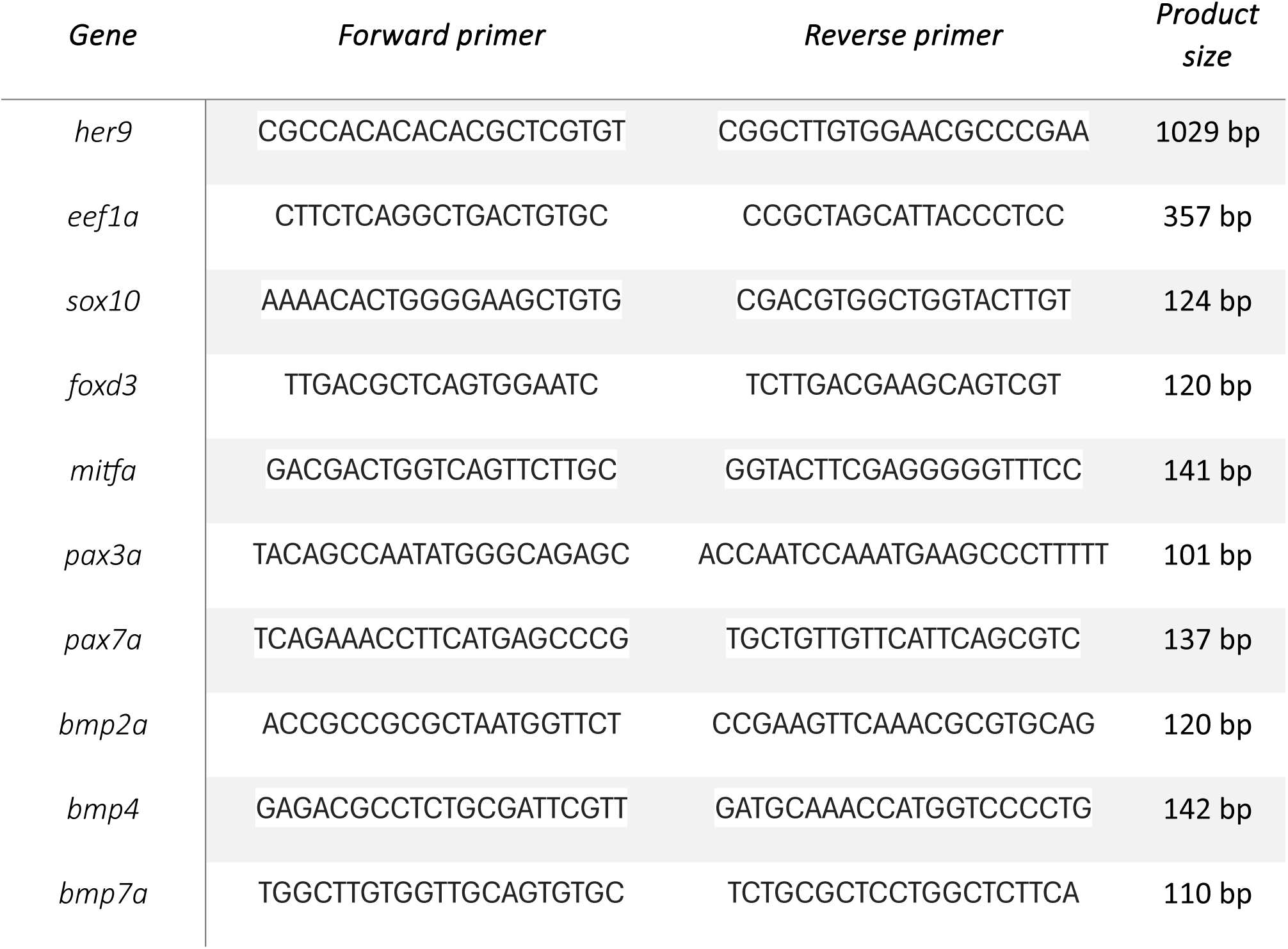
Primer sequences used in this study.

### RT-PCR and real-time quantitative RT-PCR (qPCR)

The GoScript Reverse Transcriptase System (Promega, Madison, WI) was used to synthesize the first strand cDNA from 1μg of the extracted RNA. PCR primers (Eurofins Genomics, Louisville, KY) were designed to amplify unique regions of the *her9*, *elfα*, and each neural crest specific gene (**Table 1**). Faststart Essential DNA Green Master mix (Roche) was used to perform qPCR on a Lightcycler 96 Real-Time PCR System (Roche). The relative transcript abundance was normalized to *eifα* expression as the housekeeping gene control (Wen et al., 2015), and was calculated as fold-change relative to WT siblings. RT-PCR and qPCR experiments were performed with three biological replicates (which included a minimum of 25 pooled embryos) and three technical replicates. RT-PCR was performed on a Mastercycler Pro thermocycler (Eppendorf, Westbury, NY).

### Immunohistochemistry of tissue sections

Immunohistochemistry was performed as previously described (Pillai-Kastoori et al., 2014). The following primary antibodies were used: HuC/D (mouse, 1:20, Invitrogen, Grand Island, NY), which recognizes enteric neurons; αGFP (mouse, 1:1000, Santa Cruz biotechnology); Zrf-1, which recognizes glial cells (mouse, 1:5000, Santa Cruz biotechnology). Alexa Fluor 488 goat anti-mouse, 488 goat anti-rabbit, 546 goat anti-rabbit, and 546 goat anti-mouse secondary antibodies (Molecular Probes, Invitrogen) were all used at 1:200 dilution. Nuclei were visualized by counterstaining with DAPI (1:10,000 dilution). Samples were mounted in 60% glycerol in PBS. Images were taken at 20x and 40x on an inverted fluorescent microscope and a Leica SP8 DLS confocal/digital light sheet system using a 40x or 63x objective. Ten sections from 10 embryos or larvae (1 section per individual) were analyzed on each slide and for each antibody.

### Whole-mount immunohistochemistry

Embryos were manually dechorionated at 24 hpf. Embryo tails were clipped for genotyping and the heads were fixed overnight in 4% paraformaldehyde at 4°C, then stored in 100% MeOH at -20°C until the genotypes were obtained. The tails were lysed to extract gDNA in 50mM NaOH at 96°C for 30 minutes. Genotypes were determined by RFLP analysis as described previously (Coomer et al., 2020). Heads of identical genotypes were pooled together for further processing. Samples were re-hydrated by washing once in 50% MeOH and twice in 1X PBS each for 5 minutes at room temperature. Samples were transferred to 1X PBS-TT (1% Triton-X, 1% Tween20) and washed 5 times for 5 minutes each. Samples were then incubated in freshly prepared blocking solution (1% Triton-X, 1% Tween20, 1% BSA, 5% Normal Donkey Serum, 1X PBS) for a minimum of 2 hrs with shaking at 4°C. Primary antibodies used were diluted in blocking solution as follows: Pax7 (DSHB:AB_528428, 1:20 dilution) and Sox10 (Proteintech 66786-1-1g:AB_2882131,1:200 dilution). Samples were incubated in primary antibody solution overnight with shaking at 4°C. The next day, samples were washed once in blocking solution for 10 minutes with shaking at 4°C, then four times in 1X PBS-TT for 15 minutes each at 4°C. After one wash in blocking solution, the samples were incubated overnight with shaking at 4°C in Cy5 Donkey Anti-Mouse secondary antibody diluted in blocking solution (Jackson ImmunoResearch, 715-175-150: AB_2340819, 1:250 dilution). Samples were then washed once in blocking solution and 4 times in 1X PBS-TT with shaking at 4°C. Samples were then mounted in 1% low melt agarose (Promega, V2111) in a glass bottom dish (MatTek, P35G-1.5-14-C). Z-stack images were obtained on an inverted Leica SP8 DLS confocal fluorescent microscope under the 10X objective. Pax7- and Sox10-positive cells anterior to the midbrain/hindbrain boundary were counted (two technical replicates). A minimum of five immunolabeled embryos was analyzed for each genotype and antibody.

### TUNEL Staining

Terminal deoxynucleotide transferase (TdT)-mediated dUTP nick end labeling (TUNEL) was performed on frozen retinal cryosections using the ApopTag Fluorescein Direct In Situ Apoptosis Detection Kit (Millipore, Billerica, MA). TUNEL staining was performed according to the manufacturer’s protocol. Images were taken at 20x and 40x on an inverted fluorescent microscope (Eclipse Ti-U, Nikon instruments). Ten sections were analyzed on each slide.

### H & E staining

Cryosections were processed through a series of ethanol and H2O washes; staining with Hematoxalin and Eosin was performed as described in (Trotter et al., 2009).

### Alcian blue staining

Zebrafish larvae (5-7 dpf) were fixed in 4% paraformaldehyde (PFA) at 4°C overnight. The larvae were incubated in Alcian blue solution (Alcian blue cationic dye; abcam.com) overnight at room temperature. The samples were washed in 100% ethanol (EtOH) and transferred through an ethanol series (90%, 80%, 70%) to 1X PBST. The embryos were digested in 100mg trypsin in 10 mL 30% saturated aqueous sodium borate. Specimens were bleached with Hydrogen peroxide (3% H_2_O_2_ in 1% KOH) until transparent and stored in 70% glycerol.

### Feeding Assay

To create the fluorescent tracker food, we mixed 100 mg of powdered larval feed (Gemma 75; Skretting) and 150 µL of Fluosphere carboxylate modified microspheres supplied in 2% solid solution (Invitrogen, Carlsbad, CA, USA) in 50 µL of deionized water. The mixture was combined into a paste and dried overnight, in the dark at room temperature. Once dry the flakes were stored at room temperature. The fluorescent tracker food was mixed with 10 mL of artificial fish water, and the feeding assay was conducted on 7 dpf larvae. Larvae were imaged prior to feeding and then placed in petri dishes containing food. After 10 min of feeding the embryos were screened for food in the gut and fish positive for food in the gut were isolated. The isolated fish were then imaged every 30 min for 2 hours to track the progression of the food through the digestive tract. After the feeding assay, the embryos were genotyped as described above. Five larvae were analyzed per genotype.

### Statistical analysis

Statistical analysis was performed using a student’s t-test or Chi-square test with GraphPad Prism software (www.graphpad.com). For comparing the number of neural crest cell and other phenotypic features a minimum of 10 WT and 10 mutant animals were examined (except for the whole-mount immunolabelling, which used five individuals from each genotype). All graphs present the mean ± the standard deviation (s.d.)

## Results

### Her9 mutants display craniofacial defects

While characterizing *her9* mutant visual system phenotypes (Coomer et al., 2020), we noted that the mutants displayed several defects outside the retina. These phenotypes included a lack of swim bladder (SB), craniofacial defects (CFD), curved body, and altered pigmentation (**Fig. 1A-C**). To further investigate how the loss of Her9 affects craniofacial development, we used light microscopy to document gross morphological changes within the facial structures of the *her9* mutants compared to their WT and heterozygous siblings. Among the progeny of *her9* heterozygous incrosses, 18% of the embryos displayed CFD; of those, 94% genotyped as homozygous mutants (*her9*^-/-^), 6% genotyped as *her9* heterozygotes, and none were homozygous WT (**Fig. 1H**). To determine whether the mutant craniofacial phenotype was associated with cartilage malformation, we performed Alcian blue staining on 5 dpf embryos from a *her9*^+/-^ heterozygous incross. The cartilage of the WT head skeleton contains two subdivisions, the neurocranium and viscerocranium, which include the ethmoid plate (e), trabeculae (tr), Meckel’s cartilage (m), palatoquadrate (pq), ceratobranchial (cb 1-5), and ceratohyal (ch) (**Fig. 1D**). Upon initial characterization of craniofacial defects in the *her9* homozygous mutants we observed two major phenotypic categories with respect to cb 1-5. In Phenotype 1 (60%), the ceratobrachial elements were present but appeared faint, smaller and abnormally shaped (**Fig. 1F-F’**), whereas in mutants with Phenotype 2 (20%), the ceratobranchial elements were missing (**Fig. 1G-G’**). Cranial width is often used as an indicator of craniofacial defects. To determine whether *her9* mutant embryos had smaller heads than their WT and heterozygous siblings, we measured the distance between the eyes (**Fig. 1I-I’**); we observed a decrease in the distance in the *her9* mutants compared to their wildtype siblings (**Fig. 1J**). Taken together, these data indicate that the loss of Her9 causes craniofacial defects including reduction or loss of cb1-5, and a smaller head.

**Figure 1.**
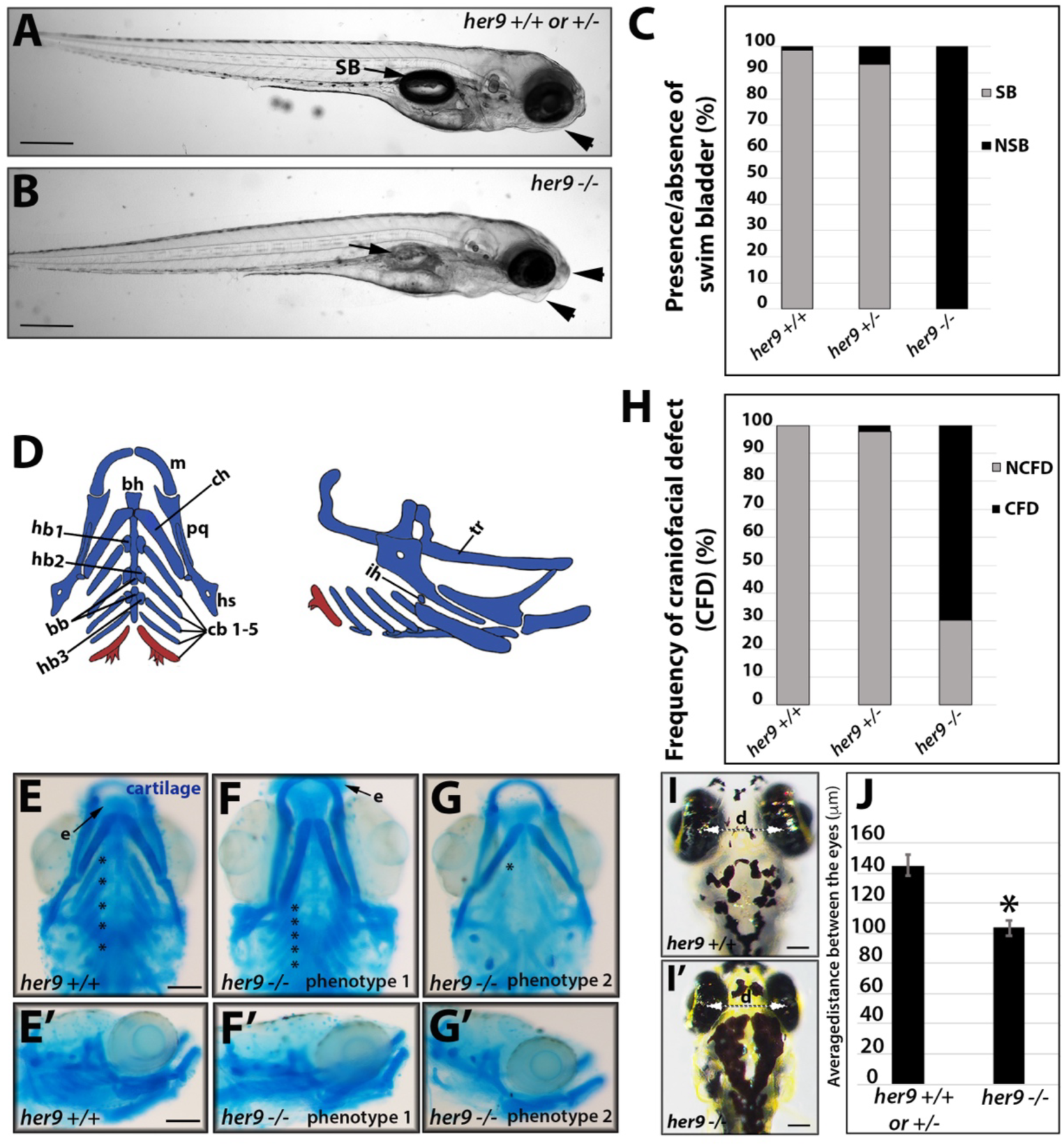
Craniofacial phenotypes in *her9* mutant zebrafish. **A-B**) Gross morphology of *her9* mutant larvae at 5 dpf compared with WT and heterozygous larvae. *Her9* mutants lack a swim bladder (SB, arrow) and have abnormally shaped jaws (arrowheads). **C**) Absence of swim bladder is a fully penetrant phenotype in *her9* mutants. **D**) Schematic of zebrafish craniofacial structures. **E-G’**) Variations in *her9* mutant craniofacial defects visualized with Alcian blue staining of cartilage. **H**) Frequency of craniofacial defects observed in *her9* vs. WT and heterozygous larvae. **I-J**) Measurement of distance between the eyes of WT and *her9* mutant larvae. Hs, hyosymplectic; bh, basihyal; ch, ceratohyal; m, Meckel’s cartilage; pq, palatoquadrate; cb, ceratobranchial arches; hb, hinge bone; tr, trabeculae; ih, interhyal; e, ethmoid plate. Scale bars = 25 μm.

### Loss of Her9 disrupts development of cranial neural crest cells

Craniofacial defects are often attributed to defects in cranial neural crest cell (CNCC) development. To determine if the craniofacial defects present in *her9* mutants are linked to the disruption of CNCC development, and at what developmental stage this phenotype is detectable, we analyzed expression of Pax7 and Sox10, which regulate NCC specification and migration, in the zebrafish head using whole mount immunohistochemistry. As early as 24 hpf, we detected a significant decrease in the number of Pax7-positive cells and a trend towards reduced Sox10-positive cells in the heads of *her9* mutant embryos when compared to their WT and heterozygous siblings (**Fig. 2A-D**). We also crossed the *her9* mutation onto the *foxd3*:GFP transgenic background; Foxd3 is a forkhead box transcription factor that is an important pre-migratory NCC gene (Plouhinec et al., 2014). At 24 hpf, we observed a decrease in the number of *foxd3:*GFP*+* cells in the *her9* mutant, specifically in the midbrain and posterior end of the hindbrain, compared to their WT and heterozygous siblings (**Sup.** Fig. 1A-B). To confirm that the changes in GFP+ cells reflected a change in endogenous *foxd3* expression, qPCR was performed; we observed a significant decrease in *foxd3* expression at 24 hpf in *her9* mutants compared to the WT embryos (**Sup.** Fig. 1C). Taken together, these results suggest that loss of Her9 alters NCC specification and migration.

**Figure 2.**
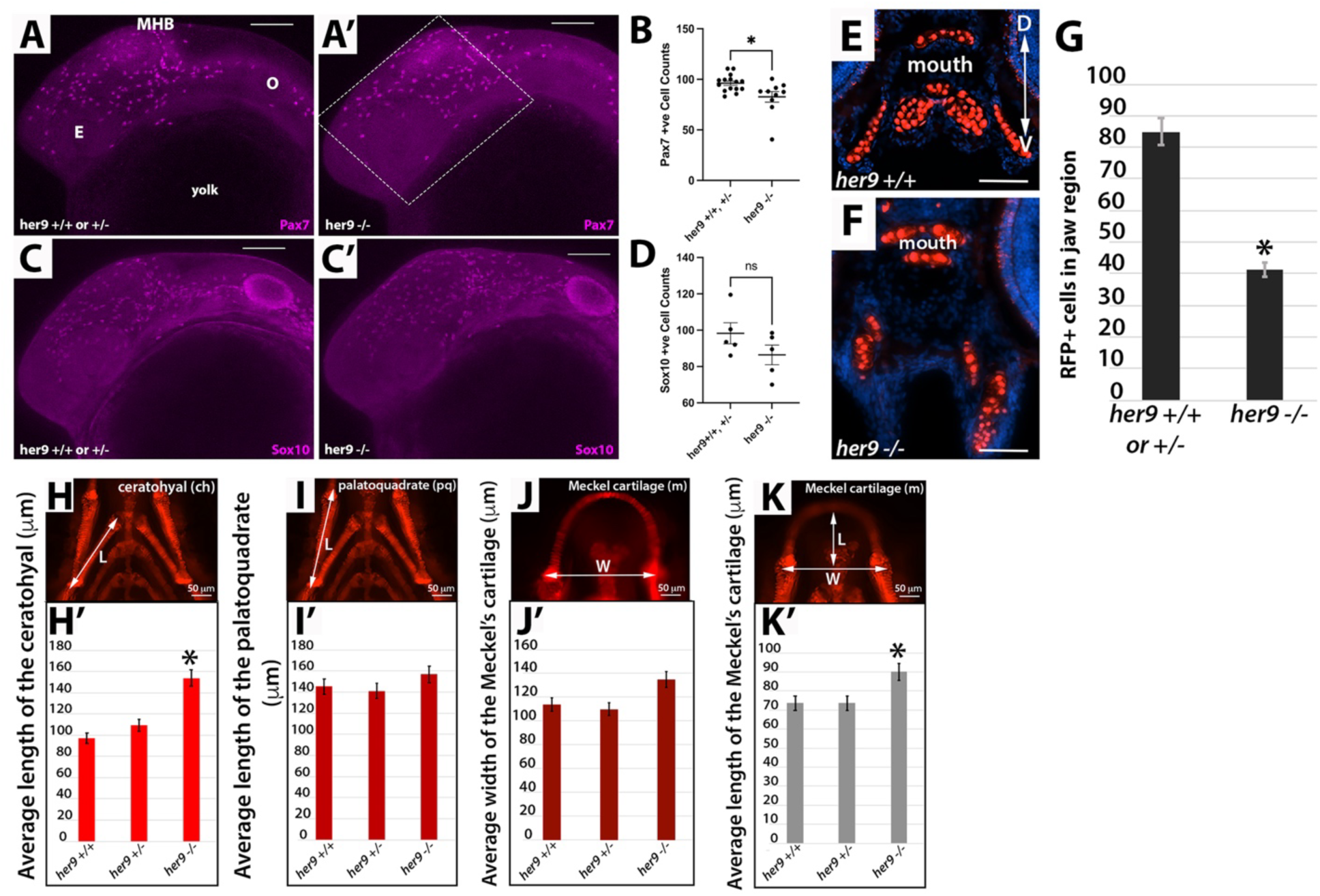
Loss of Her9 results in reduced CNCC and viserocranium structural defects. **A-D**) Whole mount immunohistochemistry for Pax7 (**A-B**) and Sox10 (**C-D**) at 24 hpf. The number of Pax7+ and Sox10+ cells anterior to the midbrain-hindbrain boundary (MHB) is reduced in *her9* mutants relative to WT and heterozygous siblings. E-G) Sox:RFP+ cells are reduced in *her9* mutant craniofacial region at 5 dpf (n=8 mutants, n=13 WT or heterozygous larvae, *p<0.001). H-K’) Analysis of craniofacial structures in WT, heterozygous, and her9 mutant larvae at 5 dpf. *p<0.05. Scale bars for A-C’, 100μm; E-F = 50 μm.

To better understand the effects of the loss of Her9 on CNCC cell development we crossed the *her9* mutation onto the *sox10*:RFP transgenic background. We then performed incrosses, collected cryosections through the heads of 5 dpf offspring and counted the number of RFP+ positive cells within the jaw/mouth region, using the optic nerve as a landmark (**Fig. 2E-G**). We observed a significant decrease in the number of RFP+ cells within the jaw/mouth area of the *her9* mutants compared to their WT and heterozygous siblings. This result suggests that the reduction in CNCC associated with loss of Her9 persists into the larval stage.

To further quantify a subset of the craniofacial defects we measured significant features of the viscerocranium. We observed a significant increase in the length of the ceratohyal in *her9* mutants compared to their WT and heterozygous siblings (**Fig. 2H-H’**). The palatoquadrate showed no significant changes in length among the different *her9* genotypes (**Fig. 2I-I’**), nor did the width of Meckel’s cartilage (**Fig. 2J-J’**). However, the length of the Meckel’s cartilage was significantly increased in the *her9* mutant larvae compared to their WT and heterozygous siblings (**Fig. 2K-K’**). We conclude from these data that the loss of Her9 disrupts the specification and/or migration of CNCC, causing a decrease in *sox10-*expressing cells in the jaw, ultimately resulting in a range of craniofacial cartilage malformations.

### Her9 mutants display pigmentation defects

To determine whether the loss of Her9 affected only the CNCC or other subtypes of NCCs, we first surveyed *her9* heterozygous embryo clutches for pigmentation defects. The pigment pattern displayed in larval zebrafish consists of three lineages derived from trunk neural crest cells (TNCC): melanophores (black), xanthophores (yellow) and iridophores (silver) (Rocha et al., 2020). These pigment subtypes are well developed by 6 dpf (**Fig. 3A-A’)**. Patterning of the melanin expressing melanophores consists of four longitudinal stripes: the dorsal and ventral stripe (extends from head to tail; black and blue arrows on **Fig. 3A-B**), the yolk stripe (extends from under the heart to the anus; red arrows on **Fig. 3A-B**), and the lateral stripe (extends through the myoseptum from somite 6 to 26; green arrows on **Figure 3A-B**) (Milos and Dingle, 1978). Although *her9* mutants possessed the same patterning as that observed in their WT and heterozygous siblings, in *her9* mutants the dorsal stripe contained fewer, but larger, pigment cells, the ventral stripe was thinner along the head and the region adjacent to the swim bladder but thicker towards the tail, and the yolk and lateral stripes appeared to have fewer melanophores (**Figs. 3A-B’ and 3I-J**). Xanthophores carry yellow pigmentation, and individual cells are hard to identify; these cells are predominantly located on the dorsal most portion of the zebrafish (**Fig. 3E-F, purple arrows**). The *her9* mutants appeared to have a darker yellow glow across the dorsal stripe compared to their WT and heterozygous siblings (**Fig. 3E-F’**). Lastly, the iridophores appear silver under ambient light and are associated with the melanophores in the dorsal, lateral, yolk stripes and the eye (**Fig. 3E-F’, asterisks**). We observed a decrease in iridophores in all these regions in the *her9* mutants relative to their WT and heterozygous siblings (**Fig. 3G-H’**).

**Figure 3.**
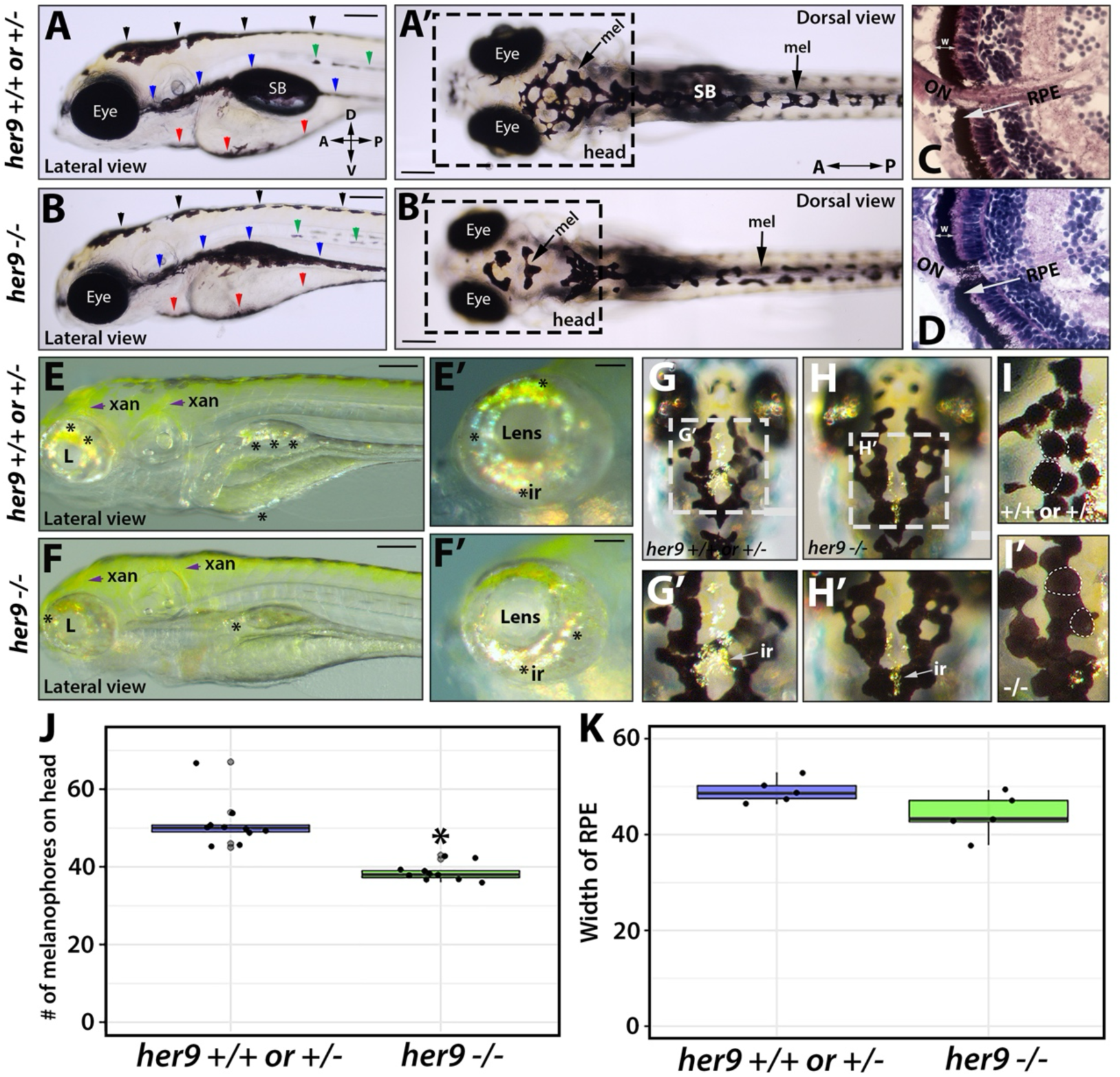
Her9 mutants display pigmentation defects. **A-B’**) Lateral (**A-B**) and dorsal (**A’-B’**) views of melanophore patterning in WT/heterozygous and *her9* mutant larvae at 6 dpf. Black arrowheads, dorsal stripe; green arrowheads, lateral stripe; blue arrowheads, ventral stripe; red arrowheads, yolk stripe; SB, swim bladder. **C-D**) H&E staining of WT/heterozygous and *her9* mutant retina at 7 dpf; the retinal pigmented epithelium (RPE) is indicated with a white arrow. **E-F’**) Xanothophore (xan) and iridophore (ir) patterning in WT/heterozygous and *her9* mutant larvae. *Her9* mutants have reduced abundance of iridiphores (asterisks) and greater spread of xanthophores (yellow) than WT or heterozygous siblings. G-I’) Dorsal view of iridiphores and melanophores on the heads of WT/heterozygous and *her9* mutant larvae. *Her9* mutants have fewer iridiphores and melanophores (quantified in **J**); **K**) there was no difference in RPE width in *her9* mutants compared to WT/heterozygous siblings. Scale bars A-B’,E, F, G, and H = 20 μm; C-D = 100 μm; E’, F’, G’, H’ and I’ = 40 μm.

The other pigmented cells in the zebrafish are the cells of the retina pigmented epithelium (RPE); however, RPE cells are not derived from NCCs. To determine whether the loss of Her9 affected all pigmented cells or just NCC-derived pigment cells, we imaged retinal sections of WT and *her9* mutant embryos at 6 dpf and observed no significant changes in the appearance or thickness of the RPE (**Fig. 3C-D, K**). We conclude that Her9 regulates the development of TNCC-derived pigment cells, but not RPE cells.

To further explore how the loss of Her9 affects pigment cell differentiation, we analyzed the expression of NCC genes associated with the three chromatophore lineages prior to, during, and after completion of their differentiation (22 and 72 hpf and 7 dpf) (Budi et al., 2011; Rocha et al., 2020). *Foxd3* expression in NCCs has been shown to drive NCCs cells towards differentiation (Petratou et al., 2018). At 48 hpf, we observed a decrease in *foxd3* expression in the *her9* mutants that was not statistically significant (**Fig. 4A**). By 72 hpf, the chromatophores begin to differentiate and all lineages are present (xanthophores, melanophores, and iridophores). Pax7 and Pax3 have both been shown regulate the xanthophore lineage, and Pax7 to inhibit melanophore differentiation (Minchin and Hughes, 2008; Miyadai et al., 2023; Nord et al., 2016). In agreement with our observation of an increase in xanthophores in the *her9* mutants, we observed an increase in the expression of both *pax3a* and *pax7a* compared to their WT and heterozygous siblings at 72 hpf (**Fig. 4B**). This increased expression persisted through 7 dpf, by which time embryonic/larval chromatophores are fully differentiated (**Fig. 4D**). Sox10, Mitfa and Foxd3 have been shown to be key players in melanophore and iridophore lineage differentiation. Sox10 regulates the expression of *mitfa,* which initiates the specification of melanocytes, while Foxd3 blocks the expression of *mitfa* and initiates the specification of iridophores (Lister et al., 1999; Petratou et al., 2018). At 72 hpf, *mitfa*, *foxd3,* and *sox10* expression were all reduced in the *her9* mutants relative to their WT and heterozygous siblings, although these reductions were not significant (**Fig. 4C**). Furthermore, by 7 dpf, *sox10* and *mitfa* expression were significantly decreased (**Fig. 4E**). Taken together, these results suggest that the loss of Her9 results in alterations in the differentiation of TNCC derived pigment cells, promoting xanthophore lineage differentiation at the expense of the melanophore and iridophore lineages.

**Figure 4.**
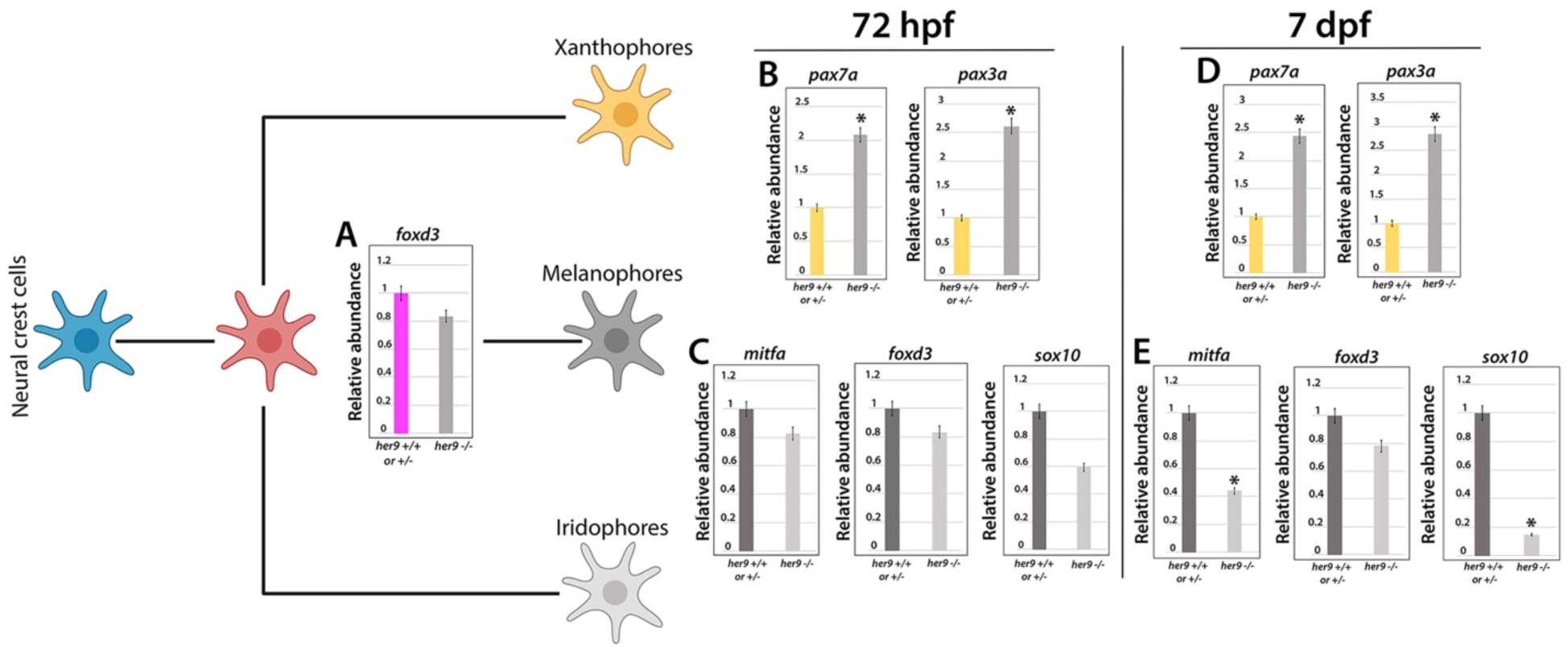
Loss of Her9 alters pigment lineage gene expression. **A**) qRT-PCR analysis of *foxd3* expression at 48 hpf. **B, D**) qRT-PCR analysis of *pax7a* and *pax3a* expression in WT/heterozygous and *her9* mutant larvae at 72 hpf and 7 dpf. **C, E**) qRT-PCR analysis of *mitfa*, *foxd3*, and *sox10* expression in WT/heterozygous and *her9* mutant larvae at 72 hpf and 7 dpf. *p<0.05

### Her9 mutant gut defects are associated with enteric nervous system abnormalities

As described in the Introduction, the enteric nervous system is derived from the vagal NCC (VNCC). Using light microscopy, we observed changes in gut morphology in the *her9* mutants, which included alterations in the shape of the gastrointestinal tract (**Sup.** Fig. 2A-D**”**), and a thicker mucosa with less epithelial folding (F) and a smaller lumen in the intestinal bulb (Ib) (**Sup.** Fig. 2D-D**”**). H&E staining on tissue sections confirmed the irregular shape and reduced average number of intestinal folds in the *her9* mutants relative to WT and heterozygous larvae **(Sup.** Fig. 2F-G). To determine if these morphological defects in the *her9* mutant gut were associated with abnormal VNCC and enteric nervous system (ENS) development, we examined transverse and sagittal sections of WT, heterozygous and mutant *her9* guts for the presence of VNCC, enteric neurons, and enteric glia at various developmental stages. First, we characterized *sox10*:RFP+ VNCC that had colonized the foregut, midgut, and hindgut at 60 hpf, before the completion of VNCC migration at 72 hpf (Uribe and Bronner, 2015) (**Fig. 5A-B”**). At this stage, we observed that some Sox10:RFP+ cells were already present in the foregut, midgut and hindgut of WT and heterozygous embryos (**Fig. 5A-A”**); however, in *her9* mutant embryos we only detected Sox10:RFP+ cells in the foregut (**Fig. 5B-B’’**), suggesting that loss of Her9 alters VNCC migration into the gut.

**Figure 5.**
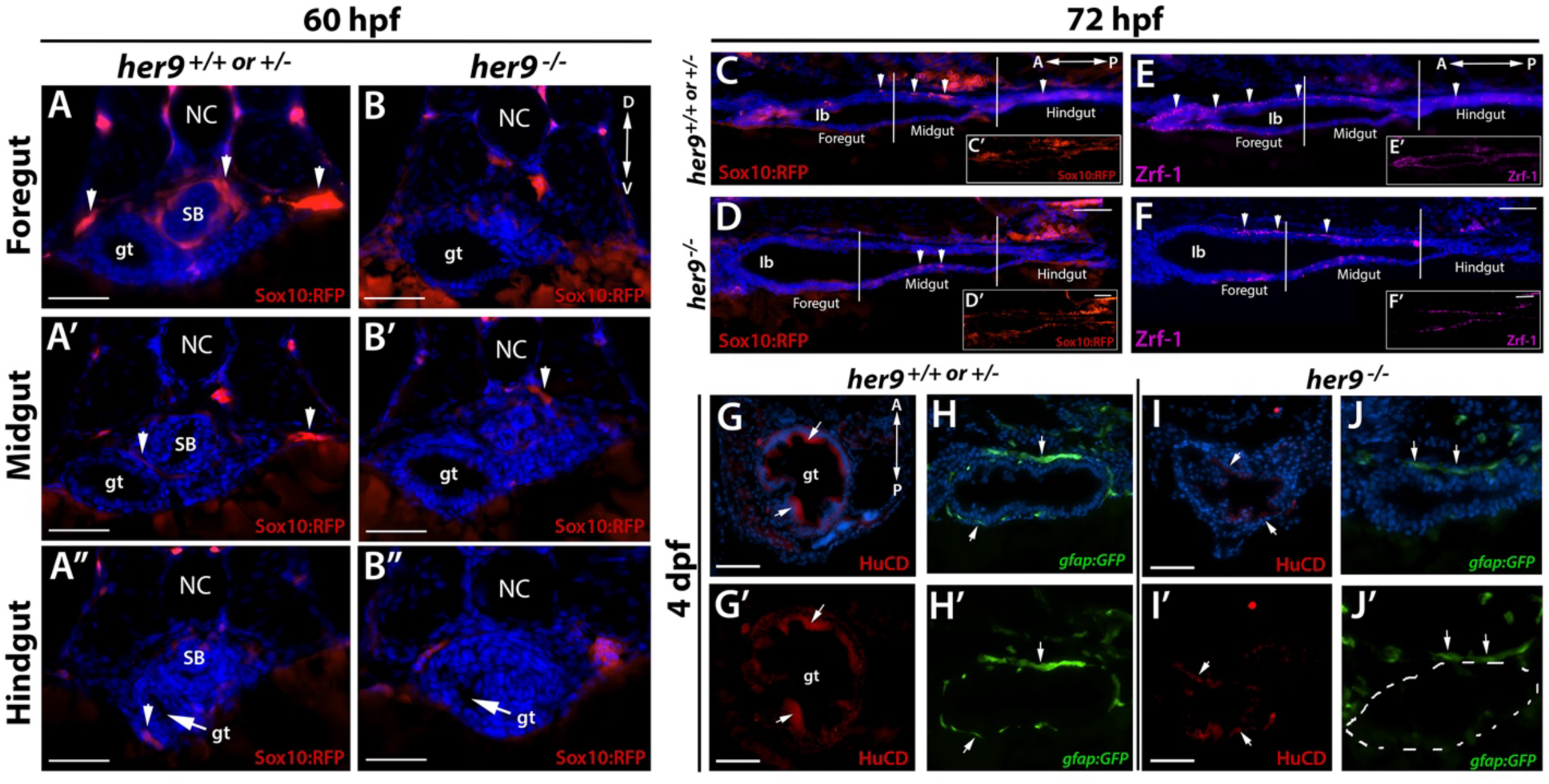
Loss of Her9 impairs NCC migration and differentiation in the gut. **A-B”**) Transverse sections through foregut, midgut, and hindgut of *sox10:RFP* WT/heterozygous and *her9* mutant embryos at 60 hpf. While some RFP+ cells have colonized all three regions of the WT/heterozygous gut (arrowheads), no RFP+ cells are visible in any region of the *her9* mutant gut at this stage. **C-D**) Lateral sections through the gastrointestinal tract of *sox10:RFP* WT/heterozygous and *her9* mutant larvae at 72 hpf. RFP+ cells are present in all three regions of the WT/heterozygous gut, but are only detected up to the midgut of *her9* mutants. E-F) Lateral sections through the gastrointestinal tract of WT/heterozygous and *her9* mutant larvae at 72 hpf immunolabeled for the glial marker Zrf-1. Zrf-1+ cells are absent from the *her9* mutant intestinal bulb (Ib) and hindgut. G-J’) Transverse sections through the gut of *gfap:GFP* WT/heterozygous and *her9* mutant larvae at 4 dpf and immunolabeled for the neuronal marker HuC/D. The *her9* mutants display fewer enteric neurons (I-I’) and fewer enteric glial cells (J-J’) than their WT/heterozygous siblings (G-G’; H-H’). SB, swim bladder; gt, gut; Ib, intestinal bulb. Scale bars A-B", G-J’ = 50 μm; C-F = 25 μm.

Examining sagittal sections at 72 hpf, when VNCC migration into the gut should be complete, we again observed Sox10:RFP+ cells in the foregut, midgut and hindgut of WT and heterozygous larvae (**Fig. 5C-C’**). However, Sox10:RFP+ cells were only observed in the foregut and midgut of the *her9* mutants (**Fig. 5D-D’**), indicating that abnormal VNCC migration persists in *her9* mutants at this stage. We used the Zrf-1 antibody, which marks GFAP-expressing glial cells, to detect enteric glia along the gastrointestinal tract. At 72 hpf, we observed a population of Zrf-1+ cells in the foregut, midgut and hindgut of the WT and heterozygous larvae (**Fig. 5E-E’**). Interestingly, although we observed Zrf-1+ cells in the foregut and midgut of the *her9* mutants, we did not observe any Zrf-1+ cells in the intestinal bulb area, and the population of Zrf1+ cells in the hindgut was reduced (**Fig. 5F-F’**). Finally, we examined cryosections of the zebrafish gut at 4 dpf to determine whether VNCC had fully differentiated into enteric neurons and glia, using the HuCD (Elav1) antibody to mark enteric neurons, and the GFAP:GFP transgenic line to label the glia. At this stage, we observed almost no HuCD+ cells in the *her9* mutant gut whereas we found several HuCD+ cells along each section of WT and heterozygous gut (**Fig. 5G-G’ and I-I’**). Likewise, we observed that the WT and heterozygous gut was populated with differentiated GFAP:GFP+ glia, while in the *her9* mutant larvae GFAP:GFP+ cells were greatly reduced (**Fig. 5H-H’ and J-J’**). Taken together, these data demonstrate that the loss of Her9 affects VNCC migration and infiltration into the gut, as well as differentiation into enteric neurons and glia.

### Her9 mutants display altered feeding habits and abnormal gut function

To further understand how the morphological and ENS defects observed in the *her9* mutant gut impacted its function, we utilized the fluorescent feeding assay described by Tang et al. (Tang et al., 2009). Larval zebrafish develop a functional intestinal tract by 5 dpf, so for our experiments we tested feeding ability at 7 dpf. We provided larvae with fluorescently labeled food, then screened them by fluorescence microscopy every 30 minutes to track the movement of food through the gut (**Fig. 6A**). Whereas approximately 10% of WT and heterozygous larvae ingested the fluorescent food within the first 10 minutes, we observed a significant decrease in the intake of food by the *her9* mutant larvae, resulting in fewer mutant larvae with food in their gut (**Fig. 6B**). The subset of larvae that displayed uptake of fluorescent food were then imaged at 60 and 90 minutes after feeding. At 60 min post feeding (mpf), we observed significantly less fluorescent signal in the gut of the *her9* mutant larvae compared to their WT and heterozygous siblings (**Fig. 6D-D’**). By 90 mpf, the fluorescent signal in the gut of WT and heterozygous larvae had expanded from the foregut into the midgut towards the hindgut (**Fig. 6E**). However, the fluorescent signal in *her9* mutants was only detectable in the foregut and the midgut, never the hindgut (**Fig. 6E’**). These results indicate that the defects in gut morphology and ENS differentiation in *her9* mutants are associated with abnormal processing of food through the gastrointestinal tract. Collectively, our results also show that loss of Her9 affects the differentiation of multiple NCC lineages, including cell types from the CNCC, TNCC, and VNCC.

**Figure 6.**
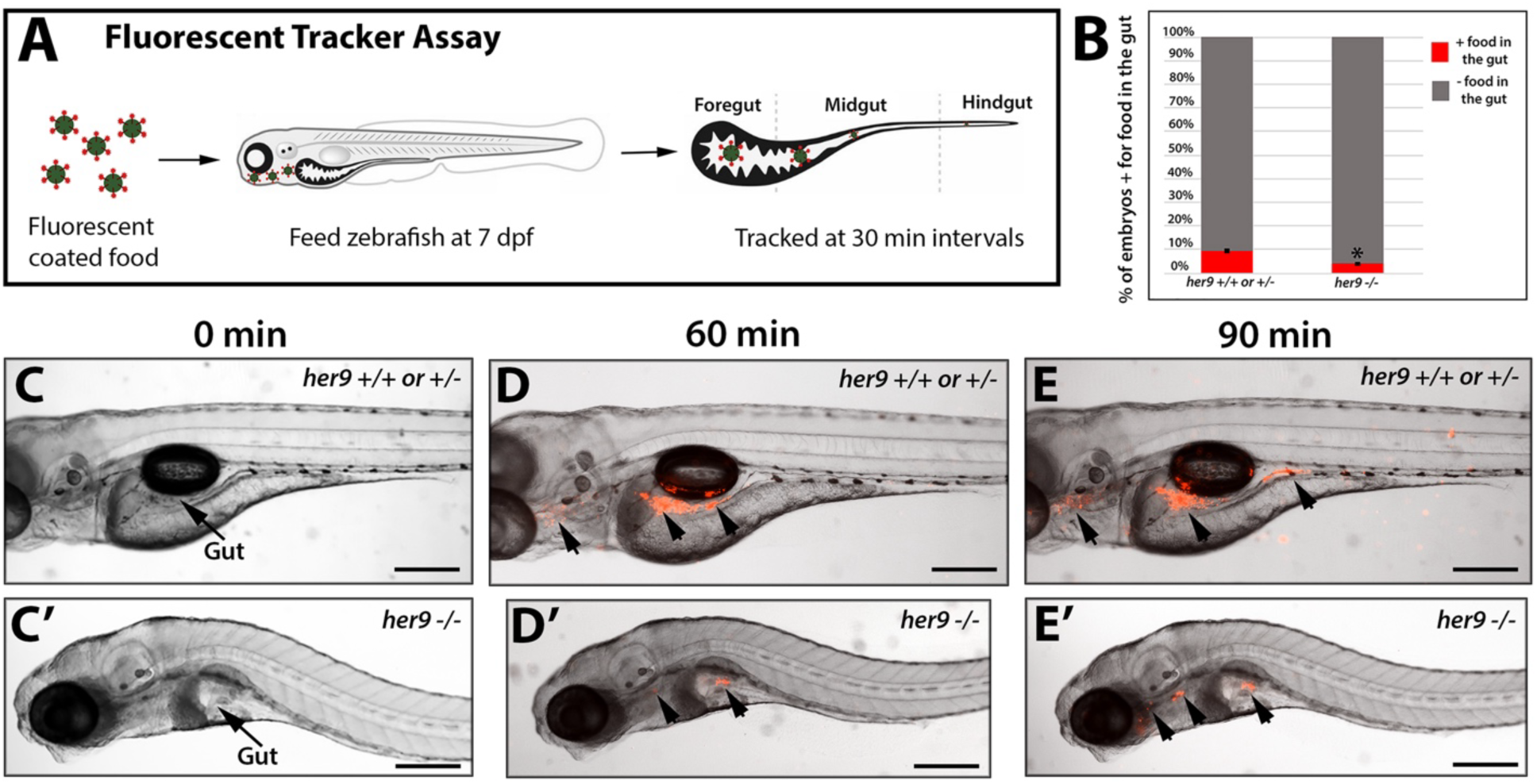
Her9 mutants display decreased food intake and abnormal transit of food through the gut. **A)** Schematic of fluorescent tracker feeding assay. **B)** Proportion of WT/heterozygous and *her9* mutant larvae that ingested fluorescently labeled food (n=15 WT/het and 9 *her9* mutants). **C-E)** Location of fluorescent food in the WT/heterozygous and *her9* mutant gut a 0, 60, and 90 minutes after feeding. By 90 minutes after feeding, the fluorescent signal had reached the hindgut of WT/heterozygous larvae, but not in the *her9* mutants. *p<0.005 Scale bars C-E’ = 20μm.

### Loss of Her9 does not disrupt dorsal root ganglia development

The dorsal root ganglia (DRG) contain peripheral sensory neurons and glia derived from trunk neural crest cells (TNCC). To determine whether the loss of Her9 affects the development of DRG neurons, we incrossed *her9* heterozygotes carrying the *sox10:RFP* transgene, and performed immunohistochemistry using the HuCD antibody to label DRG neurons at 72 hpf. We detected Sox10:RFP+ cells at the locations where DRG develops in both WT, heterozygous, and *her9* mutant larvae, although on some sections the RFP+ signal was reduced relative to WT in the *her9* mutants (**Fig. 7A, B, C-D’**). The location and labeling intensity of HuC/D also appeared similar between all three genotypes, suggesting that the DRG neurons develop normally in *her9* mutants (**Fig. 7A’, B’**). Quantification of the number of Sox10:RFP+ DRG along the body axis revealed a small decrease in the *her9* mutants relative to WT and heterozygous larvae, but it was not significant when normalized to overall body size (**Fig. 7E**). Taken together, our data indicates that Her9 is not required for development of the DRG neurons, although we cannot rule out an effect of Her9 loss on the DRG glia. This result also shows that while Her9 regulates the development of several diverse NCC lineages, it is not essential for differentiation of every NCC derived cell type.

**Figure 7.**
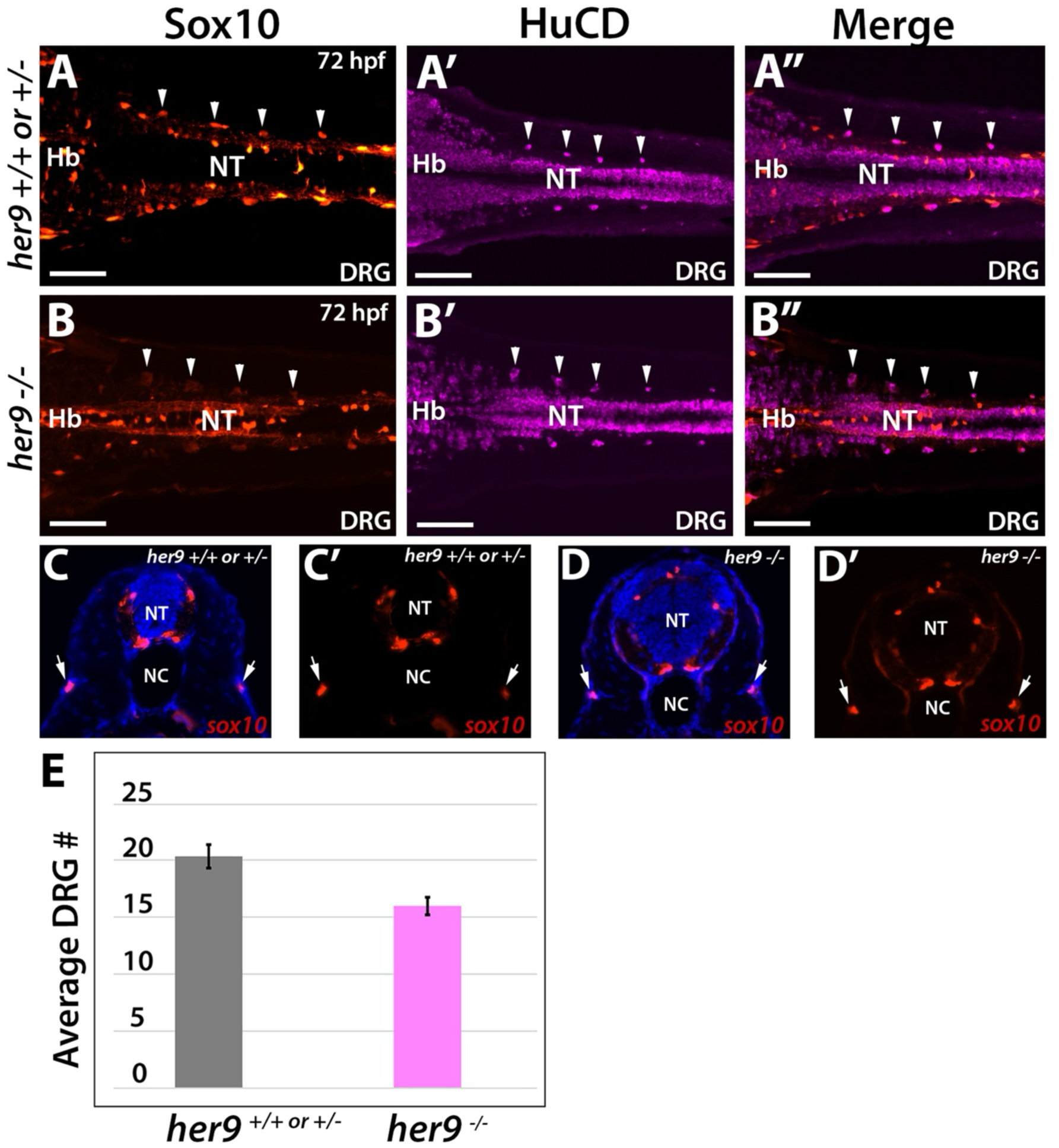
Loss of Her9 has minimal effects on dorsal root ganglia (DRG) neurons. **A-B”**) Longitudinal sections of *sox10:RFP* WT/heterozygous (**A-A”**) and *her9* mutant (**B-B”**) larvae immunolabeled with the neuronal marker HuC/D. No differences were observed between in the number or distribution of RFP+ and HuCD+ DRG neurons (arrowheads). **C-D’)** Sox10+ DRG neurons were also detected in transverse sections of *sox10:RFP* WT/heterozygous (**C-C’**) and *her9* mutant (**D-D’**) larvae. **E)** Quantification of the average number of DRG neurons per genotype; averages were not significantly different. NT, neural tube; NC, notochord; Hb, hindbrain.) Scale bars = 20 μm.

### Loss of Her9 leads to death of CNCC and VNCC derived cells

Disruption in the expression of several NCC genes, such as *foxd3*, *bmp* ligands, and *vgll2a*, has been shown to affect the survival of NCCs (Huang et al., 2016; Johnson et al., 2011; Stewart et al., 2006). To determine whether the loss of Her9 caused NCC death, we performed TUNEL staining on cryosections from 5 and 7 dpf larvae. We did not observe any cell death in the *her9* mutant pigment cells (not shown), but we did observe a significant increase in TUNEL+ cells in the *her9* mutant mouth and jaw compared to the WT and heterozygous larvae at 5 dpf (**Fig. 8A-B’**). We also observed a significant increase in the number TUNEL+ cells in the gut of *her9* mutants compared to the WT and heterozygous larvae at 7 dpf (**Fig. 8C-D’**). These results suggest that Her9 is required for the survival of specific NCC derivatives.

**Figure 8.**
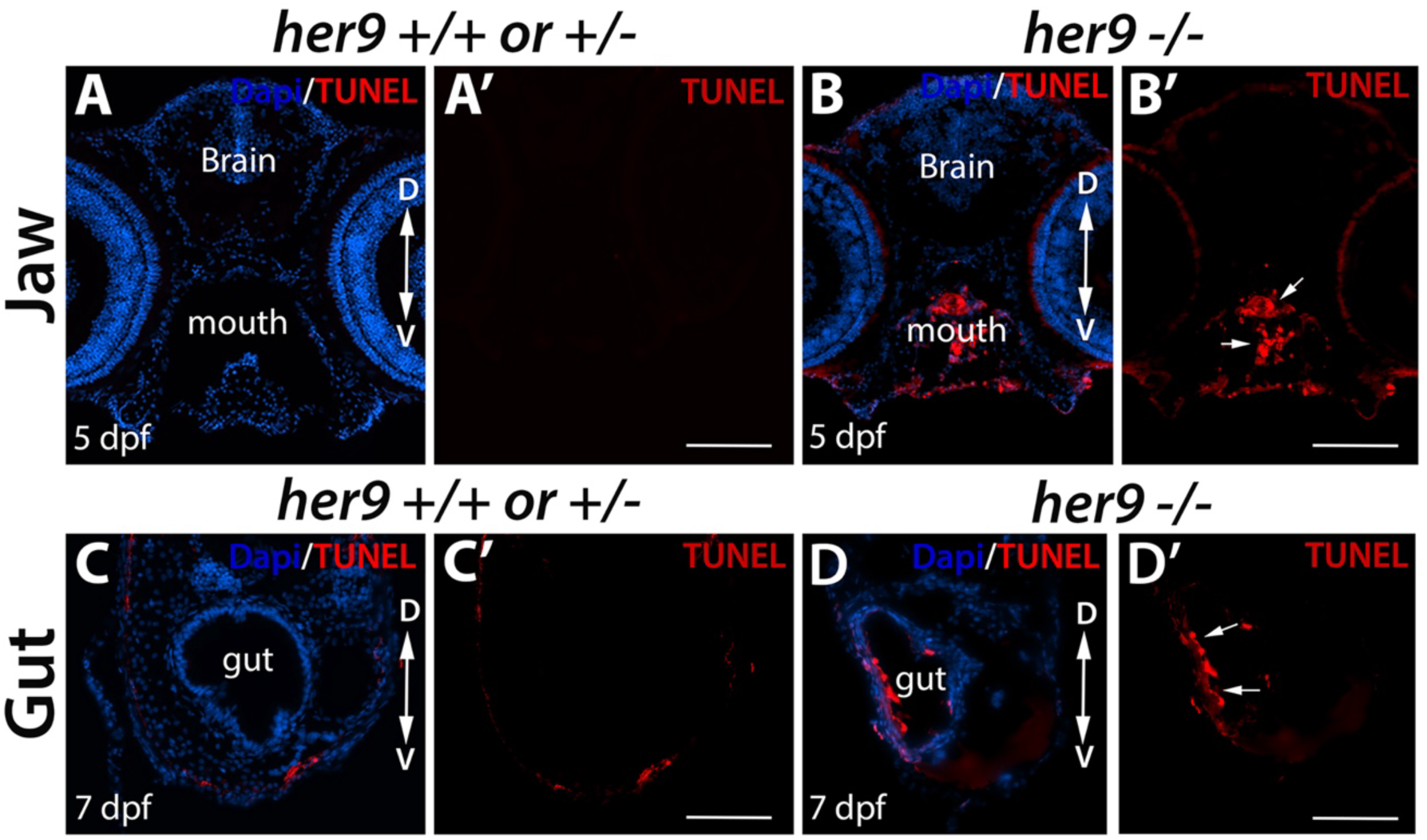
Loss of Her9 results in apoptosis of CNCC- and VNCC-derived cell types. **A-B’**) TUNEL staining of sections through the heads of 5 dpf WT/heterozygous (**A-A’**) and *her9* mutant (**B-B’**) larvae. TUNEL+ cells are detectable in the mouth and jaw region of *her9* mutants (arrows). **C-D’**) TUNEL staining of sections through the gut of 7 dpf WT/heterozygous (**C-C’**) and *her9* mutant (**D-D’**) larvae. TUNEL+ cells are detectable in the gut of *her9* mutants (arrows). Scale bars A-B’ = 20 μm; C-D’ = 50 μm.

### Her9 mutants have an altered Bmp gene expression during significant phases of NCC development

Finally, to begin addressing the mechanism by which loss of Her9 might lead to such widespread NCC phenotypes, we examined the expression of BMP ligands at various developmental stages. BMP signaling is known to regulate NCC induction, migration and differentiation, along with other signaling pathways (such as FGF, Wnt, and Hh) (Schumacher et al., 2011). We used qPCR to analyze the expression of *bmp2a*, *bmp4* and *bmp7* at 24, 48, 72 hpf and 5 dpf in WT/hets and *her9* mutant embryos and larvae. At 24 hpf, when NCC cell induction is complete and CNCC migration is in progress, we observed a decrease in the expression of all BMP ligands in *her9* mutants relative to their WT and heterozygous siblings (**Fig. 9A**). However, by 48 hpf, when NCC are migrating, BMP ligand expression was increased in the *her9* mutants relative to WT and heterozygous embryos (**Fig. 9B**). This increase in BMP ligand gene expression persisted at 72 hpf (**Fig. 9C**), when some NCC are still migrating and others are differentiating, and at 5 dpf (**Fig. 9D**), by which time most NCC derivatives have completed migration and differentiation. These data suggest that loss of Her9 causes an upregulation in BMP ligand expression and BMP signaling following NCC induction. They also indicate that upregulation of BMP signaling upon loss of Her9 may perturb NCC migration, differentiation, and survival.

**Figure 9.**
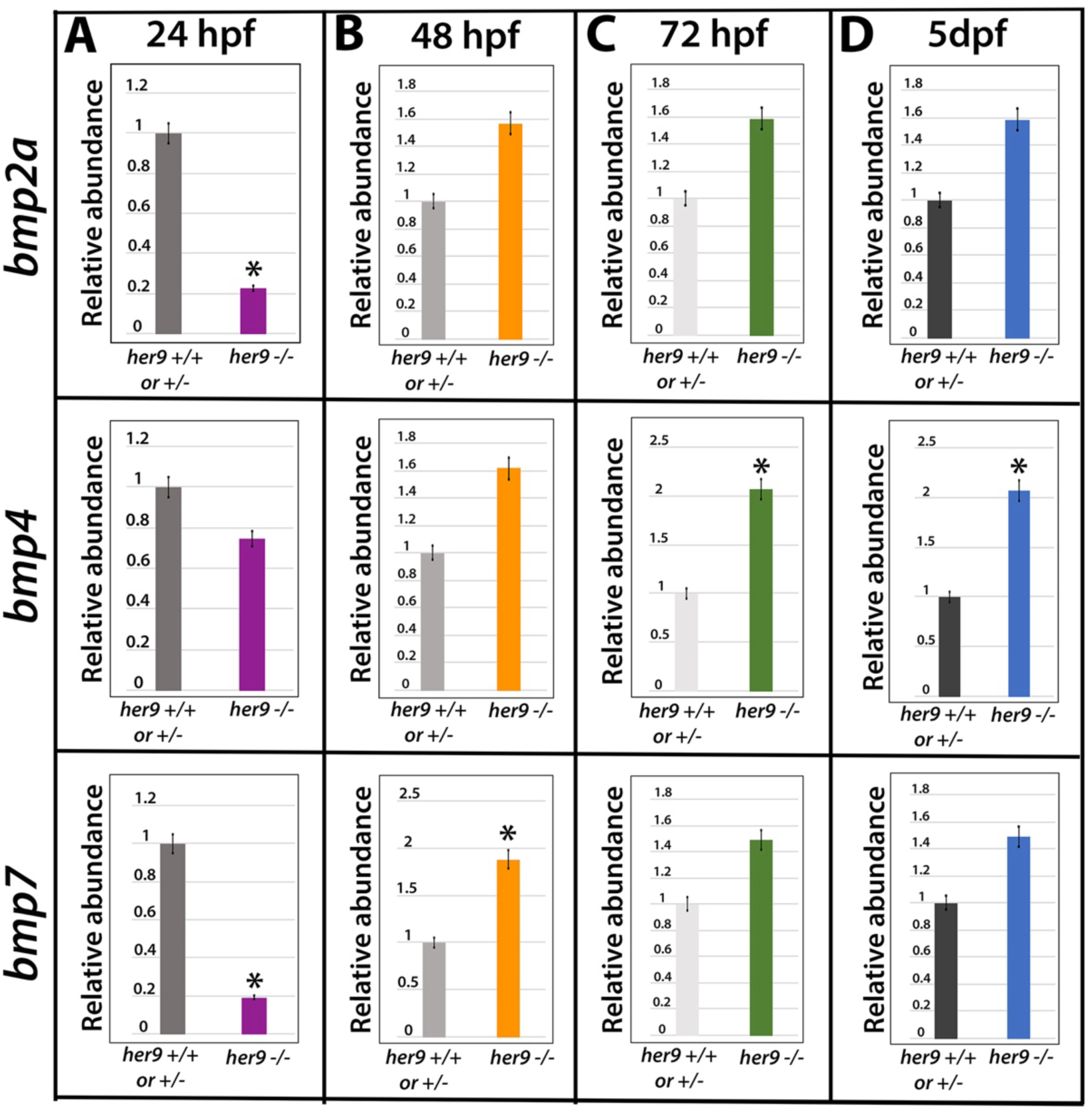
Expression of Bmp ligand genes is upregulated in *her9* mutant larvae. qRT-PCR for *bmp2a*, *bmp4*, and *bmp7* expression in whole embryos/larvae at 24 hpf (**A**), 48 hpf (**B**), 72 hpf (**C**), and 5 dpf (**D**). At 24 hpf, expression of *bmp2a* and *bmp7* is decreased in *her9* mutants relative to WT/heterozygous embryos. However, at later time points (48 and 72 hpf, 5 dpf), expression of *bmp2a*, *bmp4*, and *bmp7* is higher in *her9* mutants than WT/heterozygous larvae. *p<0.05.

## Discussion

In this study, we analyzed the NCC phenotypes present in the zebrafish *her9* genetic mutant. Her9/Hes4 is a bHLH-O transcription repressor and, like many other members of the Hairy/Hes family, is known to be involved in the regulation of neurogenesis. Here, we show that functional Her9 is also required for the migration, differentiation, and survival of specific NCC derivatives. Moreover, our results suggest that Her9 functions upstream of essential genes within NCC gene regulatory network (GRN) such as *bmp2a/4/7*, *foxd3* and *sox10*.

### Her9/Hes4 during early NCC development

Our analysis of the *her9* mutant indicates that loss of Her9 may affect NCC induction and specification, but it is likely not essential, because we do observe *sox10+* or *foxd3+* NCC and differentiated NCC-derived structures in *her9* mutants. Previous studies have shown that morpholino-mediated knockdown of *her9/hes4* results in reduced expression of NCC genes (*slug and foxd3*), leading to the decrease or depletion of specific NCC subpopulations, such as dorsal root ganglia and sympathetic and enteric neurons while melanocyte number was delayed but not significantly affected (Stewart et al., 2006). In our study, we found that *her9* mutants presented with significant reduction of *foxd3* expression at 24 hpf, a reduced number of *foxd3:*GFP positive cells, and gross morphological craniofacial and gut defects. The significant decrease in *foxd3* expression and decrease in the number of *foxd3* expressing cells could indicate a decrease in the pre-migratory NCC population. Additionally, the decreased expression of *sox10* and decreased number of *sox10* expressing cells in CNCC and ENS NCC populations indicate that there were fewer cells specified, implying Her9 is involved in induction and specification of NCCs.

Bone morphogenetic proteins (BMP) have been shown to act in coordination with Noggin to create a gradient that allows for the specification of NCC at a specific concentration (Marchant et al., 1998). At high levels BMP activity induces epidermis, intermediate levels of BMP induce NCCs, and the lack of BMP is required for neural ectoderm formation (Schumacher et al., 2011). In our *her9* mutant, there is a slight decrease in the expression of BMP ligands at 24 hpf, followed by a significant increase in BMP ligand expression at later timepoints. The relatively modest change in BMP ligand expression during the period of NCC induction and specification may explain why we did not observe a complete lack of *foxd3+* and *sox10+* NCC at this stage.

### Her9 is necessary for the migration and differentiation of NCC lineages

When CNCC fail to properly migrate and differentiate, morphological defects in the face, neck and cardiovascular system arise (Hutson and Kirby, 2007; Tobin et al., 2008). The *her9* mutants display multiple craniofacial defects including missing pharyngeal arches, cleft palate and protruding mandible. CNCC migration begins with directed migration from the hindbrain along the dorsolateral pathway towards the branchial arch and finally entering and invading those arches (Kulesa et al., 2010). Improper development of branchial pouches in the *her9* mutants is therefore indicative of defects in NCC migration. We did not observe any obvious morphological cardiac defects in *her9* mutants, but further investigation of specific cardiac structures is necessary to rigorously investigate effects of *her9* on heart development.

Moving from CNCC to other NCC lineages, we observed significant changes in pigment cell differentiation in the *her9* mutants, which included a decrease in the number of melanophores and iridophores accompanied by an increase in xanthophores, suggesting a possible fate switch in the pigment cell lineage. These changes were associated with a sustained increase in the expression of NCC genes *pax3* and *pax7* from 3-7 dpf. In the developing gut, we observed reduced numbers of VNCC-derived neurons and glia, as well as delayed migration from the foregut to the hindgut. These alterations were associated with reduced food passage through the gut, phenotypes which mimic the neurocristopathy Hirschsprung’s disease. Taken together, these data show that, in addition to the CNCC, Her9 is required for the proper development of TNCC and VNCC, and provide supporting evidence that loss of Her9 disrupts not only NCC induction or specification, but also migration and differentiation of NCC derivatives.

One NCC-derived tissue that did not appear to be affected by loss of Her9 is the DRG. DRG have been shown to be downstream of Wnt and Hh signaling (Ungos et al., 2003; Won et al., 2011) but not BMP signaling during NCC development; this could explain why we did not see any significant effects on this NCC subpopulation. Although we did not observe significant differences in DRG neurons by IHC with the HuCD antibody, it remains to be determined whether other cell types (such as satellite glia) in the DRG are altered upon loss of Her9.

### Her9 and NCC survival

Previously, Xhairy2/Hes4 was shown to be required for the proliferation and survival of NCCs (Nagatomo and Hashimoto, 2007). Our studies support a role for Her9 in NCC survival, as we observed a significant increase in cell death in the pharyngeal arches and ENS of *her9* mutants at 5 dpf. These phenotypes could be a result of the misexpression of transcription factors within the NCC GRN which have dual roles in cell development and survival. For example, disruption in the expression of *foxd3* in mice leads to the death of embryonic stem cells (Hanna et al., 2002). Loss of *foxd3* expression also leads to cell death in a subpopulation of NCC in the hindbrain that contributes to neurons, glia, and pharyngeal arches (Stewart et al., 2006). Therefore, the consistent downregulation of *foxd3* in the *her9* mutants could contribute to the significant cell death we observed in the cranial NCC and the neurons and glia of the ENS.

## Conclusion

In this study, we describe an array of NCC phenotypes in *her9* mutant zebrafish. In a previous study, we documented retinal phenotypes associated with loss of Her9. Whether the defects in the retina and in NCC derivatives are both caused by dysregulation of BMP signaling remains to be determined. Currently, most eye or retinal phenotypes associated with neurocristopathies are characterized by morphological and structural changes rather than cellular defects in the retina. This could very well be due to the extensive nature of some of the phenotypes in neurocristopathy patients, causing retinal cellular phenotypes to be overlooked and difficult to characterize. It is also possible that Her9 is playing multiple roles through separate pathways, one in the retina and another in NCCs. Further exploration into the pathways that influence *her9* expression during NCC and retinal development would clarify some of these remaining questions and possibly provide key insight into less characterized retinal defects that could link them to neurocristopathies.

## Acknowledgments

The authors would like to thank Lucas Vieira Francisco, Brandi Bolton, and Deep Patel for excellent zebrafish care. We also thank Dr. Jakub Famulski (University of Kentucky) for generously providing some of the zebrafish transgenic lines.

---

**Supplemental Figure 1.**
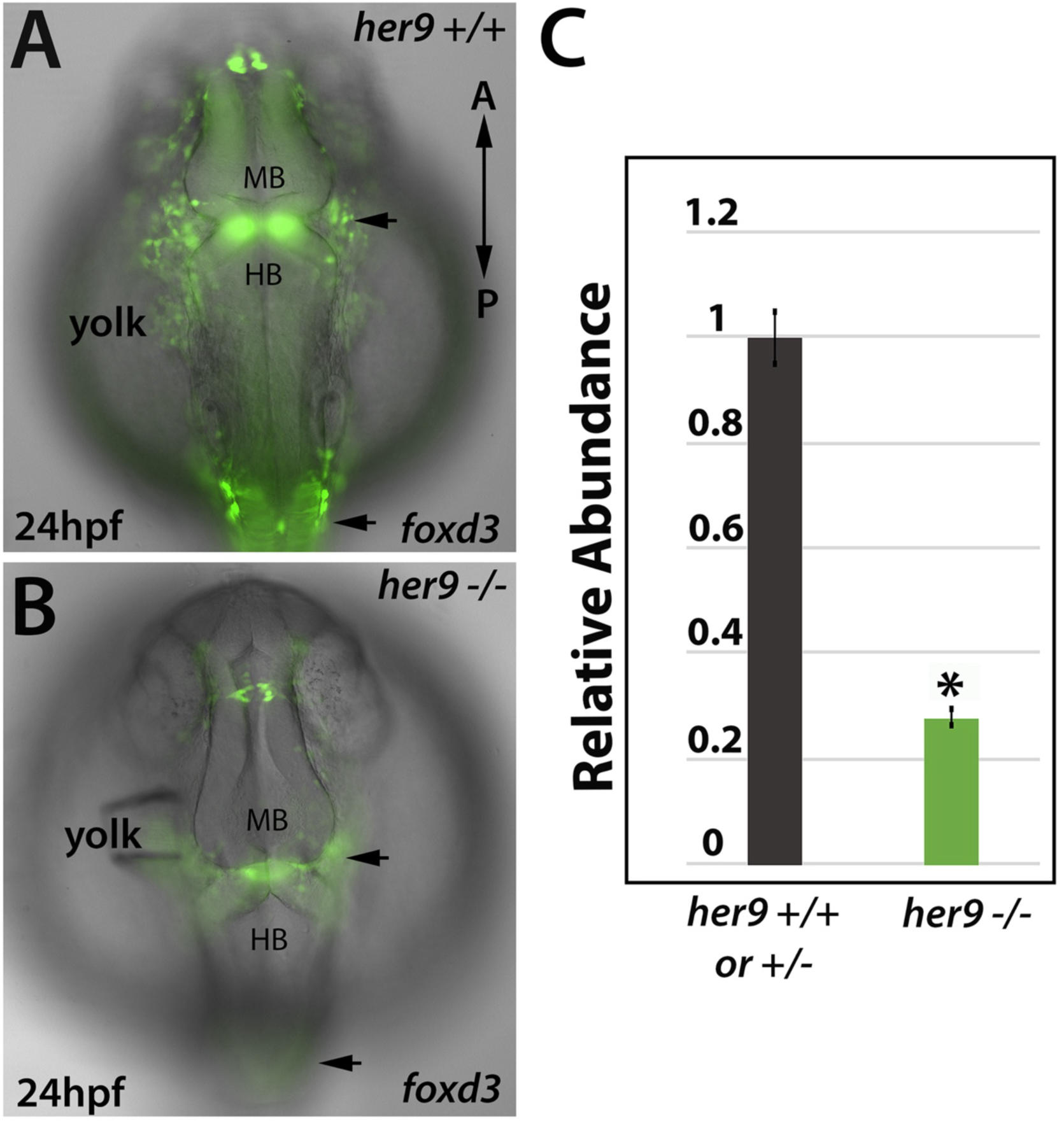
Foxd3 expression is reduced in *her9* mutants. **A-B)** Dorsal images of 24 hpf WT (A) and *her9* mutant (B) embryos crossed onto the *foxd3*:GFP transgenic background. In *her9* mutants, GFP reporter expression is diminished relative to WT in the midbrain (MB), hindbrain (HB), and the developing branchial pouches (top arrows). **C)** qRT-PCR for *foxd3* expression at 24 hpf showed a significant decrease in the *her9* mutant embryos compared to WT/heterozygous siblings. Scale bars = 20 μm.

**Supplemental Figure 2.**
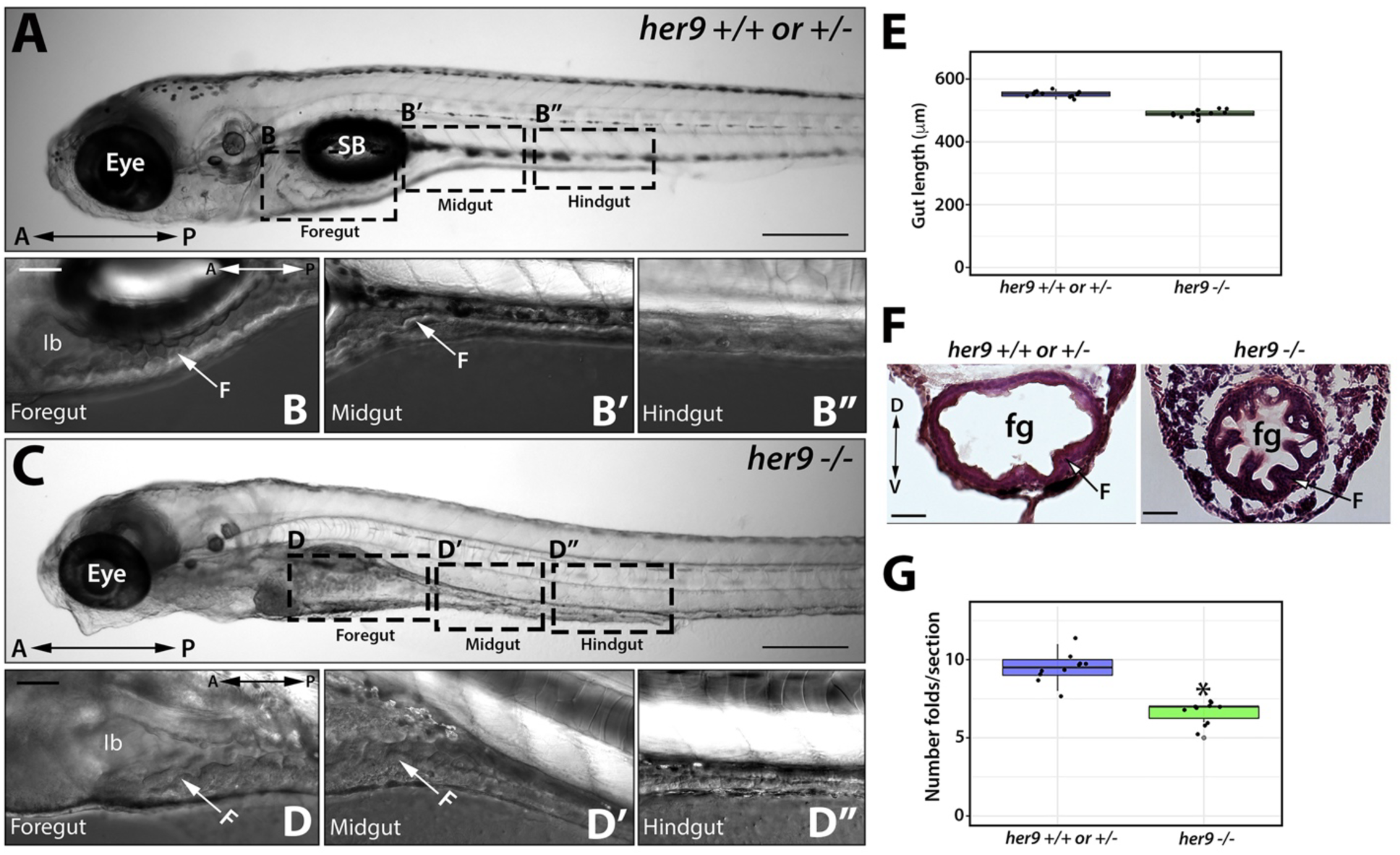
Her9 mutants display abnormal gut morphology. **A-D”)** Bright field images of 5 dpf WT/heterozygous (**A-B”**) and *her9* mutant (**C-D”**) larvae, with 40X magnification of the foregut (**B, D**), midgut (**B’, D’**) and hindgut (**B”, D”**). Compared to the WT/heterozygous images, the *her9* mutant gut appears irregularly shaped and with fewer folds (F). **E)** The length of the gut was not significantly different between WT/heterozygous and *her9* mutant larvae. **F**) Transverse H&E stained sections through the foregut of WT/heterozygous and *her9* mutant larvae. The *her9* mutant foregut displayed a smaller lumen and abormal folds compared with WT/heterozygous foregut. **G**) The average number of intestinal folds per section was reduced in *her9* mutants relative to WT/heterozygous siblings. SB, swim bladder; Ib, intestinal bulb; F, intestinal folds. Scale bars A, C = 20 μm; B-B" and F = 50 μm.

